# Tactile Stimulation Designs Adapted to Clinical Settings Result in Reliable fMRI-based Somatosensory Digit Maps

**DOI:** 10.1101/2022.12.23.521816

**Authors:** Till Steinbach, Judith Eck, Inge Timmers, Emma Biggs, Rainer Goebel, Renate Schweizer, Amanda Kaas

**Affiliations:** Department of Cognitive Neuroscience, Maastricht University, Maastricht, The Netherlands; Brain Innovation B.V., Maastricht, The Netherlands; Department of Medical and Clinical Psychology, Tilburg University, Tilburg, The Netherlands; Department of Anesthesiology, Perioperative and Pain Medicine, Stanford University School of Medicine, Palo Alto, California, USA; Functional Imaging Laboratory, German Primate Center, Göttingen, Germany; Leibniz ScienceCampus Primate Cognition, Göttingen, Germany

**Keywords:** fMRI, human, somatosensory cortex, tactile stimulation, retest reliability, translational

## Abstract

A wide range of neurological diseases with impaired motor functioning of the upper extremities are accompanied by impairments of somatosensory functioning, which are often undescribed but can provide crucial information for diagnostics, treatment selection, and follow-up. Therefore, a reliable description of the functional representation of the digits in the somatosensory cortex would be a highly valuable, but currently lacking, tool in the clinical context. Task-based functional Magnetic Resonance Imaging of passive tactile stimulation provides an indirect, but valid description of the layout of the digit map in the primary somatosensory cortex. However, to fulfill the specific requirements for clinical application, the presently established approaches need to be adapted and subsequently assessed for feasibility and retest reliability, in order to provide informative parameters for the description of the evoked digit activations. Accordingly, the present high-field 3T fMRI study compares the performance of two established digit mapping designs - travelling wave (TW) and blocked design (BD) - for passive tactile stimulation of the five digits, adapted to reduce the time requirements to just below 15 minutes. To be able to assess the retest reliability unaffected by any clinical conditions, the study was performed on neurotypical participants. The results show that both stimulation designs evoke significant and distinct activation clusters in the primary somatosensory cortex of all participants for all five digits. The average spatial locations of the center of gravities across participants show the common succession of distinct digit representation along the central sulcus. The cortical extent elicited activation, which is generally larger for the thumb and the index finger, also shows comparable average values across the two approaches. Less overlap of activation between neighboring digits was obtained in BD, consistent with the distinct single digit neuronal representations. A high retest reliability was obtained for the location of the digit activation, displaying stable center of gravity locations across sessions for both stimulation designs. This is contrasted by only medium to low retest reliability for the extent and overlap of the digit activations, indicating discrepancies across sessions. These results demonstrate the capacity of shortened fMRI digit mapping approaches (both TW and BD) to obtain the full layout of single digit cortical activations on the level of the individual, which together with the high reliability of the location of the digit representation over time indicates both approaches are clinically applicable.

## 1. INTRODUCTION

The clinical assessment of somatosensory functioning in neurological diseases with motor deficits, especially in stroke, is standard practice with physicians, physiotherapists, and occupational therapists for general prognosis, to assist treatment planning and to review progress of impaired motor functions of the limbs (Pumpa et al., 2015; Winward et al., 1999). The clinical relevance of this practice is confirmed by estimations that somatosensory impairments are present in roughly every second stroke patient, with numbers possibly being underestimated due to a lack of application of standardized somatosensory testing procedures (Kessner et al., 2016; Sullivan & Hedman, 2008). As the motor system fundamentally relies on the somatosensory system, providing the feedback for effective movement control, disturbances in somatosensation will result in impairments (Borich et al., 2015), and need to be treated alongside motor disturbances.

The study of neuroplasticity and reorganization in the primary somatosensory cortex (SI) has a long history, with separate neuronal representations of the five fingers already described in the somatosensory homunculus (Penfield & Boldrey, 1937), and later confirmed and specified by electrophysiological recordings in monkeys (Iwamura et al., 1983; Merzenich et al., 1987). Merzenich and colleagues (1983) studied these cortical maps in response to digit amputation and found substantial translocations of the discontinuities between digits, i.e., significant map reorganizations, providing strong evidence that the cortical digit maps are dynamically maintained. In a separate line of investigations, comparing the specifics of the detailed digit map outline across individual adult monkeys, they could furthermore show distinguishable variations within the general digit topography. The authors described these individual map layouts as a consequence of the difference in the typical use of the hand and deduced a lifelong adaptability of the digit map (Merzenich et al., 1987).

Reorganization and neuroplasticity of the somatosensory cortical digit maps has also been illustrated in humans using non-invasive neuromagnetic source imaging (i.e., magnetoencephalography (MEG)) (Elbert et al., 1995; Flor et al., 1995). Specifically, reorganization due to omission of somatosensory input has been demonstrated in upper limb amputees. Tactile stimulation of fingers of the unaffected hand and lip resulted in shorter distances between the cortical lip representations and the mirrored digit representations (from the contralateral SI) in the affected hemisphere implying reorganization. The extent of reorganization was furthermore correlated with the occurrence of phantom limb pain (Flor et al., 1995). The adaptability of the digit map, i.e., the effect of lifelong specific usage of the fingers on the individual maps, has been demonstrated in professional string players. The musicians showed a larger spread of cortical digit representations of their extensively used fingers of the left hand as compared to the digit representations of the left, non-dominant hand of control subjects (Elbert et al., 1995). These two studies exemplify the numerous research describing the reorganization of the somatosensory system associated with loss of sensory input due to injury in the periphery or the central nervous system (Flor et al., 2006) as well as the adaptability of the digit map under the conditions of various long- and short-term changes in the usage of the hands (Braun et al., 2000; Kolasinski, Makin, Logan, et al., 2016; Recanzone et al., 1992; Sterr et al., 1998; but also see Andersen & Aflalo, 2022). These findings were readily substantiated in early clinically related MEG studies describing reorganization, e.g. enlarged extension of the entire digit representation due to mono-hemispheric lesions (Rossini et al., 1998) as well as recovery-induced changes in stroke patients showing a correlation of an increase in amplitude of the first positive defection (P1m) in the somatosensory evoked magnetic field with a better 2-point-discrimination ability, indicating the neurophysiological basis for the recovery of discriminative touch (Wikström et al., 2000).

Despite these examples, and the small but growing number of clinical studies investigating the role of the somatosensory cortex in stroke or lesion related recovery of somatosensory and motor functions (Kessner et al., 2016; S. Meyer et al., 2014; Zandvliet et al., 2020), there are still substantial challenges for the comprehensive and effective implementation of somatosensory based interventions. These challenges are primarily the standardization of somatosensory assessments (Carey et al., 2016; Pumpa et al., 2015) and the development of evidence-based treatments of (e.g.,) stroke patients (Carey et al., 2011), requiring an improved understanding of the changes in SI related to functional outcomes (Sullivan & Hedman, 2008), and how somatosensory intervention strategies can impact SI (Borich et al., 2015; Carey et al., 2016). An essential first step towards these goals is the realization of standardized assessments to gain insight into SI reorganization.

An important tool in studying injury- or usage-related reorganization is through descriptions of the digit topography, which can be achieved with functional magnetic resonance imaging (fMRI). In earlier studies, blood oxygen level dependent (BOLD)-fMRI has been established as a neuroimaging technique capable of capturing the sequential representations of the five digits of one hand in the contralateral primary somatosensory cortex (Kurth et al., 2000; Maldjian et al., 1999; Nelson & Chen, 2008; Schweizer et al., 2008). With the availability of ultra-high field 7T MRI in combination with multi-channel coils and advanced MRI measurement techniques, even higher spatial and temporal resolution results in more detailed maps within the primary somatosensory digit area and beyond (Arbuckle et al., 2022; Huber et al., 2020; Sanchez-Panchuelo et al., 2010). The specific topic of reorganization and phantom limb pain is one example where these developments have already broadened and advanced the field (Makin & Flor, 2020).

These advances in our understanding of the digit topography in SI can be highly valuable when applied to preclinical and even clinical research questions. However, to achieve the translation of these research-focused digit mapping approaches to the clinic, a compromise between state-of-the-art and the reality of the clinical environment has to be considered. Specifically, this means a return from ultra-high field 7T MRI to the more readily available 3T MRI, accompanied by the challenge of decreased spatial resolution which is not always best suited for the narrow width of SI (Fischl & Dale, 2000; J. R. Meyer et al., 1996), potentially resulting in partial volume effects. Furthermore, considering the burden on patients, mapping procedures have to be as short as possible, but still long enough to reliably achieve a full digit map in the individual. Also, passive tactile stimulation is typically preferred over a motor task, such as button presses, due to potential limitations in mobility and/or possible constraints to follow the instructions of an active task. In addition to the feasibility of obtaining the digit topography in clinical conditions, high retest reliability is essential. This is necessary to accurately capture dynamic reorganization over time, as a biomarker of changes in the usage of the hand and fingers and hence recovery.

To address this need for a clinically appropriate digit mapping protocol, which will allow us to study the mechanisms needed to develop evidence-based screening and treatment, the present study focuses on: (i) the feasibility of mapping the somatosensory digit representations within the parameter range needed for clinical use and (ii) identifying the retest reliability of digit mapping for these parameters. To achieve these objectives, two 3T fMRI sessions were conducted, applying two different mapping approaches, with durations of less than 15 minutes, during which (passive) tactile stimulation was applied to the fingertips of all five digits of the right hand. The two mapping approaches applied, *Travelling Wave* (TW) and *Blocked Design* (BD), have both been successfully implemented to obtain high spatial resolution single digit maps in the somatosensory cortex by sequential stimulation of the five digits (Besle et al., 2013; Kolasinski, Makin, Jbabdi, et al., 2016; Sanchez-Panchuelo et al., 2010; Schweisfurth et al., 2014, 2018), the difference laying in shorter (TW) vs. longer (BD) stimulus periods and the absence (TW) or presence (BD) of non-stimulation periods. The study was conducted in neurotypical volunteers. This allowed the description of the retest reliability with the least number of confounding factors influencing the layout of the digit map in the second, quasi-identical, MRI session, which would not have been possible with patients due to possible recovery processes. To characterize the achieved fMRI-based digit maps within Brodmann Area 3b of SI, measures of the location and the area of single digit activations were obtained, as well as overlap of neighboring digit activations. To reflect the envisioned usage in the clinical context, not only the group averages and their variance of all measures are reported, but also the values of the individual participants describing the entire range of the data. This mimics the situation of the clinical application, in which the obtained measures are on the level of the individual, being the starting point for diagnostics and the stratification for treatment approaches.

## 2. METHODS AND MATERIALS

### 2.1. Participants

Seven neurotypical participants (mean age: 24.9 ± 2.1 years; four females, three males) were recruited via convenience sampling. All participants were right-handed, with no self-reported injuries or sensory impairments to their hands, and no self-reported history of neurological disorders. Participants received €30 for their participation. The experiment was approved by the local ethics review committee of Maastricht University and participants gave written informed consent about their participation in the experiment.

### 2.2. Study procedure

In this within-subject design, participants attended two MRI sessions (separated by 14.3 ± 7.4 days; Range: 7-24 days). Between both sessions, participants continued their regular activities, with no specific instructions given. Each two-hour-session consisted of one structural MR acquisition and four functional digit mapping measurements. Eight additional functional MR measurements obtained in each session are not considered in the present analysis. These other measurements included mapping of the non-dominant hand, both hands at the same time, and active finger tapping sequences. Since the focus of the present analysis is on retest reliability, both sessions included the same measurements.

### 2.3. Mapping Procedure

During the functional measurements, participants were asked to keep their eyes open and to fixate on a cross back-projected to the center of a screen at the end of the scanner bore. At the same time passive vibrotactile stimulation was applied through 5 separate modules of a mini piezotactile stimulator (mPTS; Dancer Design, Merseyside, United Kingdom; *Fig. 1B*) attached to the most distal phalanx of the five digits of the right hand. A stimulation frequency of 25 Hz was delivered through a metal probe (diameter: 6 mm) positioned centrally at the top of the module, moving approximately 0.5 mm up and down. This frequency corresponds to the flutter range, optimally activating the rapidly adapting type 1 afferent fibers (Saal & Bensmaia, 2014). Order and timing of the stimulation was defined by two different stimulation designs, the traveling wave (TW) and the blocked design (BD), as specified below.

**Figure 1.**
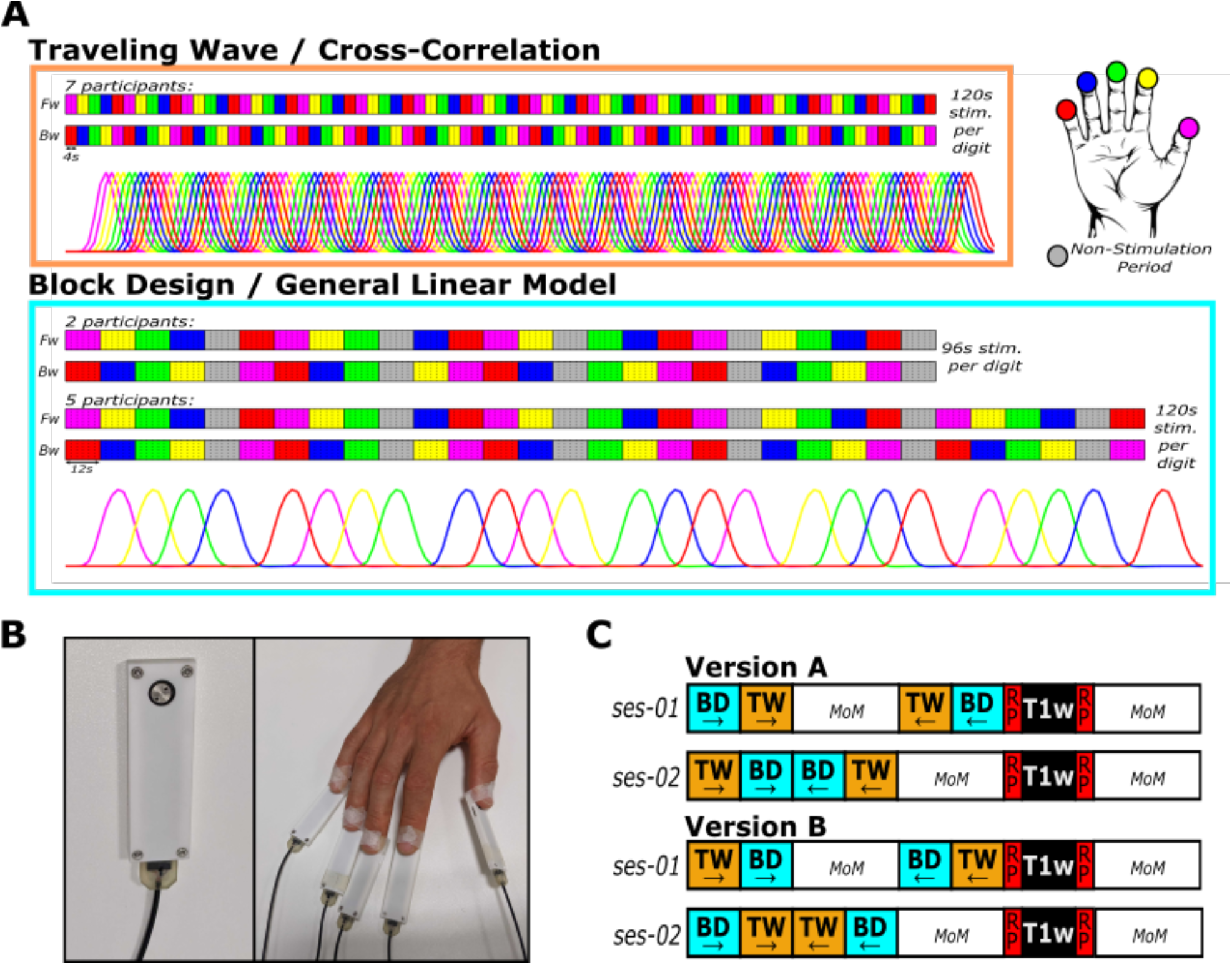
Stimulation device and stimulation designs. **A**. Traveling wave (TW) and blocked design (BD) consist of two stimulation directions: Forward (Fw; upper row) with stimulation moving from D1 to D5 and Backward (Bw; lower row; D5 to D1). The TW consists of 4s stimulations of each digit which is repeated 15 times over both stimulation directions. The BD consists of 12s stimulations of each digit and a 12s non-stimulation rest period (gray) after every fourth digit stimulation. TW was analyzed using a cross correlational analysis consisting of two models per digit shifted by one TR (2s). BD was analyzed using a standard GLM analysis with one predictor per digit. **B**. The mini piezotactile stimulator (mPTS) was used to deliver 25Hz vibrotactile stimulation to the most distal phalanx of all five digits of the right hand through the round metal probe on each module (left panel). The modules are attached to the participant’s fingers (right panel). **C**. The order of the stimulation designs within the two sessions was counterbalanced across the participants by assigning them to either Version A or B. In both versions, both sessions began with the Fw runs of the two designs, starting with either BD or TW (counterbalanced across sessions). Then, the Bw runs were acquired in the opposite order. During the first session, multiple other measures not relevant for the current experiment (MoM) were acquired between the Fw and Bw runs. In the second session, MoM were acquired after the Bw runs. Lastly, the T1-weighted scans (T1w) were always preceded and followed by functional scans with reversed phase encoding direction (RP), required for EPI distortion correction of the functional scans of interest. All sessions ended with the acquisitions of MoM.

#### 2.3.1. Traveling Wave

During the measurement runs with TW stimulation (*Fig. 1A*), the five digits of the right hand were repeatedly stimulated in successive anatomical order: thumb (D1), index finger (D2), middle finger (D3), ring finger (D4), little finger (D5). Two different directions of stimulation were acquired in separate measurement runs: forward with successive stimulation from D1 to D5 and backward with successive stimulation from D5 to D1.

For one digit, the stimulation period lasted four seconds, resulting in a complete TW-cycles of 20 s for all five digits. Each forward or backward TW fMRI measurement run consisted of 15 cycles, resulting in a total acquisition time of 5:20 minutes, including 10 s non-stimulation periods at the beginning and the end. The combined acquisition time of both TW stimulation directions was 10:40 minutes, i.e., each digit was stimulated for 120 s during each stimulation run.

#### 2.3.2. Blocked Design

The BD (*Fig. 1A*) was based on the approach used by Schweizer and colleagues (Schweisfurth et al., 2014, 2015, 2018). Digits were stimulated in anatomical succession (from D1 to D5), non-stimulation rest periods were introduced after every fourth stimulation, so that the position of the rest changes within the sequential stimulation of the five digits. This results in a longer BD cycle in which each digit is stimulated four times during one cycle and the five rest periods occur once before and once after the stimulation of each of the five digits (D1, D2, D3, D4, rest, D5, D1, D2, D3, rest, D4, D5, D1, D2, rest, D3, D4, D5, D1, rest, D2, D3, D4, D5, rest). Not only the included ‘wandering’ rest period, also the length of the stimulation period of 12 s, is more similar to a classical block design. The forward (D1-D5) as well as the backward (D5-D1) stimulation direction consisted of one BD cycle lasting 300 sec.

The BD runs of the first two participants were matched in acquisition time to the TW: one BD cycle plus a non-stimulation period of 10 s at the start and end, resulting in an acquisition time of 5:20 minutes and a combined acquisition time of 10:40 minutes for both stimulation directions. This resulted in 96 s total stimulation time per digit compared to 120 s total stimulation time per digit in the TW design.

For the remaining five participants, it was decided to keep the total stimulation duration for each finger constant across BD and TW design, instead of the total acquisition time. Therefore, one 12 s stimulation per digit was added to the end of each stimulation direction run of the BD, resulting in a total stimulation duration of 120 s per digit, which is identical to the stimulation duration in the TW design. Additionally, a rest period was added after the fourth stimulation. This version of the BD, thus, included one BD cycle (300 s) plus five digit stimulations and one rest period (72 s in total) as well as a non-stimulation period of 8 s at the begin of each run and 20 s at the end, resulting in a total acquisition time of 6:40 minutes per run and a combined acquisition time of 13:20 minutes for both stimulation directions.

The two stimulation directions of both mapping designs were acquired consecutively in a counterbalanced fashion (*Fig. 1C*).

#### 2.3.3. Attentional Task

Since attention to touch increases somatosensory cortical activation (Johansen-Berg et al., 2000) and counteracts potential habituation effects, short time segments without stimulation were inserted within the period of the vibrotactile stimulation (Besle et al., 2013; Schweisfurth et al., 2011, 2014, 2015, 2018). In the TW design, seven 100 ms long disruptions were included in every digit stimulation spaced by 500 ms. Since both the length of the digit stimulation and the disruption were rather short and difficult to perceive, participants were asked to count the total number of cycles to assure that their attention was focused on the stimulation.

In the BD, 150 ms long disruptions were included within the stimulation of each digit. The timing and number within a stimulation period was pseudorandomized and identical per digit, but different between digits. That is, stimulation of D1 and D5 always included five disruptions, D2 and D3 both included six disruptions, and D4 included four disruptions. Participants were asked to count the total number of disruptions in the vibrotactile stimulation during each run of the BD (Schweisfurth et al., 2011, 2014, 2015, 2018).

### 2.4. Data Acquisition

MRI data were obtained at a Siemens 3T Prisma Fit system with a 64-channel head coil (Siemens Healthcare, Erlangen, Germany). Anatomical T1-weighted data were acquired with the 3D whole brain coverage Alzheimer’s Disease Neuroimaging Initiative (ADNI) Magnetization Prepared Rapid Gradient Echo (MPRage) sequence (TR = 2300 ms; TE = 2.98 ms; flip angle = 9°; Bandwidth = 240 Hz/Px; FoV = 256 × 256 mm; number of slices = 192; spatial resolution = 1 × 1 × 1 mm3). Functional data were acquired using T2*-weighted Gradient Echo Echo Planar Imaging (GE-EPI) (TR = 2000 ms; TE = 30 ms; flip angle = 77°; Bandwidth = 1786 Hz/Px; Multiband Acceleration Factor = 2; GRAPPA Factor = 2; FoV = 200 × 200 mm; number of slices = 64; spatial resolution = 2 × 2 × 2 mm3). An additional functional measurement of two volumes with reversed phase encoding direction (posterior to anterior) was recorded before and after the structural measurement (*Fig. 1C*), later to be used to correct for EPI distortions (Breman et al., 2020). These reversed phase encoding data were, due to technical difficulties, not available for two participants, so no EPI distortion correction could be performed for these two participants.

To assure participants’ comfort during the MR-measurements, as well as to minimize head motion, foam padding was used inside the head coil. Additionally, medical tape was attached from one side of the head coil across the participants’ forehead to the other side of the head coil. With this, any head movement resulted in a slight pull on the participant’s skin, providing detailed tactile feedback to the participant and hence further reducing any head motion during data acquisition (Krause et al., 2019).

### 2.5. Data Analysis

MRI analysis was carried out using BrainVoyager (Version 22.0, Brain Innovation, Maastricht, The Netherlands), NeuroElf toolbox (Version 1.1) for Matlab (Version 2020a, MathWorks), and in-house Matlab and Python (Version 3.7.4) scripts.

#### 2.5.1. Data Preprocessing

Structural MR images were intensity inhomogeneity corrected, transformed into MNI space (ICBM-MNI 152) and for each participant averaged across the two sessions. Functional MRI data were slice-time corrected (cubic spline interpolation), as well as motion corrected and aligned (detection: trilinear; correction: sinc interpolation) to the run and volume that was closest to the anatomical scan during each session. To correct for low-frequency noise, the functional data were high-pass filtered with a cut-off frequency of 0.01 Hz. EPI distortion correction was obtained with the BrainVoyager plugin COPE 1.1 and to preserve the 2 mm isotropic spatial resolution, no extra spatial smoothing was performed.

Coregistration of functional MRI data to within-session structural MRI scans was achieved with boundary-based registration and transformation of functional data into MNI space was performed with sinc interpolation; in the same step, the spatial resolution of the functional data was interpolated to 1 mm isotropic resolution.

#### 2.5.2. Definition of the Region of Interest

The region of interest (ROI) was defined following the procedure described by Valente and colleagues (Valente et al., 2019), taking inter-individual anatomical differences into account without the need to manually draw ROIs for each participant.

Individual MNI-normalized cortical surface meshes of the left-hemispheric white-gray matter boundary were created and aligned using cortex based alignment (CBA) in BrainVoyager (Fischl et al., 1999; Goebel et al., 2006). An earlier study has shown a significant and meaningful improvement of the macro-anatomical correspondence of the primary somatosensory hand region between participants after CBA as compared to a volumetric normalization approach (Frost & Goebel, 2012). For CBA, an individual curvature map is created from each participant’s hemisphere, with all maps being subsequently aligned to a dynamic group average, representing the average curvature information of the specific sample. The cortex-based aligned and averaged cortical mesh of the seven participants was used to draw a ROI manually, covering the anterior wall of the postcentral gyrus opposite of the hand knob area (Yousry, 1997) stretching from the fundus of the central sulcus towards the crown of the postcentral gyrus (see *Fig. 2*). This ROI is expected to cover the hand area of somatosensory Brodmann area (BA) 3b as well as parts of BA 3a and BA 1, while reducing the risk of including the large blood vessels in BA1 (Frigeri et al., 2015). This ROI was subsequently back projected to each participant’s cortical surface mesh using the information from the CBA procedure and in a second step sampled from the surface space to the volume space by expanding it by 2 mm both towards the white matter and the cerebrospinal fluid to assure coverage of the entire gray matter ribbon in that region.

**Figure 2.**
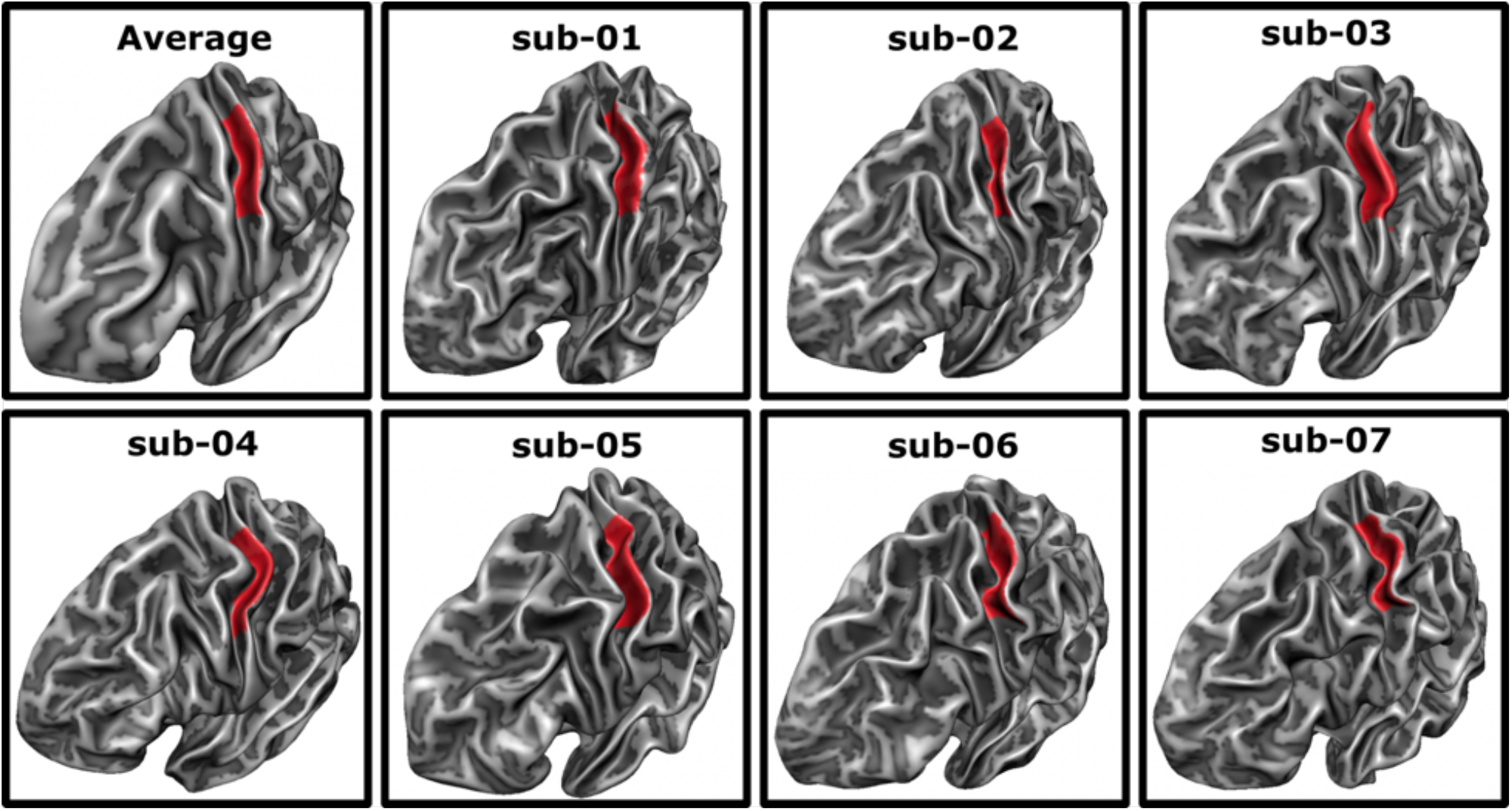
Average and individual regions of interest. The region of interest (ROI) was defined on the averaged surface mesh (top left) as the posterior bank of the central sulcus opposite to the motor hand area. Subsequently, it was backprojected to each participant’s individual surface mesh, covering a similar area in all participants.

#### 2.5.3. First Level Analyses

##### 2.5.3.1. Cross-Correlational Analysis of the Traveling Wave Design

The TW was analyzed based on the cross-correlation approach outlined by Kolasinski and colleagues (Kolasinski, Makin, Jbabdi, et al., 2016) and originally used in retinotopy (Engel et al., 1997). A reference predictor was created using a boxcar function based on the single digit stimulation length of 4 s on and 16 s off (stimulation of the other 4 digits), repeated 15 times, reflecting the fifteen TW cycles. The on-periods of the predictor were then iteratively shifted by 2 s (i.e., the TR of the functional scans) creating a total of 10 predictors, to reflect activity across the entire cycle. To account for the hemodynamic delay each predictor was convolved with a two-gamma canonical response function.

Next, the time courses of all voxels within the ROI were correlated with all 10 time-shifted predictors. To allow for statistical tests of the resulting correlation coefficients and correct for their non-normal sampling, Fisher z-transformation was applied (Silver & Dunlap, 1987). Each predictor was then assigned to one finger (i.e., the finger that was stimulated during the boxcar predictor’s on-period), thus two sets of predictors were assigned to each finger as each single stimulation duration (4 s) was double the functional TR (2 s; *Fig. 1A*). The Fisher z-transformed correlation values of the two models corresponding to the same finger were then averaged for each voxel. Then for each voxel within the ROI the correlation values from the forward and backward run were averaged, resulting in five statistical maps of the somatosensory region. Values in these maps that exceeded a false discovery rate (FDR) corrected threshold (based on all voxels within the ROI) of q(FDR) < .05 were labeled as active.

##### 2.5.3.2. General Linear Model Analysis of the Blocked Design

For the analysis of the BD, a general linear model (GLM) analysis was used within the ROI. Each finger was modeled by a boxcar regressor which was convolved with a two-gamma hemodynamic response to model the hemodynamic response function (HRF; *Fig. 1A*). A fixed effects analysis including both the forward and backward run was used for each participant. Each finger regressor was contrasted against the mean of the remaining four digit regressors. Voxels that exceeded an FDR corrected threshold of q(FDR) > .05 were labeled as active in the somatosensory ROI.

##### 2.5.3.3. Functional Vein Artifacts

To control for the influence of draining veins on the BOLD signal, voxels that showed significant activity for three or more digits (Schweisfurth et al., 2011) were excluded from further analysis, both for TW and BD. This was done separately per session and design; thus, a voxel that was excluded in one session or design was not necessarily excluded in the other session or design. The rationale behind this is that in a normal clinical setting, data from only one design and one session will be available and therefore should be sufficient to exclude all potential draining vein artifacts.

##### 2.5.3.4. Visual Inspection

To visually inspect the somatosensory activation maps that were elicited by the passive vibrotactile stimulation of the participants’ digits, the data of each session and design were projected to the flattened cortical surface representation of each participant’s left hemisphere. The functional data were projected onto the flattened surface using trilinear interpolation. Data were sampled from a 4 mm ribbon around the reconstructed white-gray matter boundary covering approximately 1 mm into the white and 3 mm into the gray matter direction, using only the maximum value in this range. This was done solely for the purpose of visualization; all analyses were performed in volume space.

##### 2.5.3.5. Definition of Activation Clusters

The activation of each digit in each design and session was defined as its largest cluster of significant BOLD-activation in volume space that contained the peak voxel (i.e., the voxel with the highest statistical value) within the ROI. In case the largest activation cluster did not contain the peak voxel, either the largest cluster or the cluster containing the peak voxel was chosen, depending on which cluster was closest to the clusters chosen for the neighboring digits.

#### 2.5.4. Description of Map Parameters

To investigate the feasibility of mapping the somatosensory digit representation using the two adapted designs (TW and BD), several parameters were extracted from the activation clusters of the first session and compared across the two designs. These parameters included two parameters concerned with the location of the digit representation (center of gravity and D1-D5 distance) and two parameters concerned with the area of activation (volume of activation and activation overlap between neighboring digits).

##### 2.5.4.1. Center of Gravity

The center of activation for each digit was defined as the center of gravity (CoG) of that digit’s activation cluster. The coordinates of the CoG were determined based on the cluster’s average coordinates weighted by each voxel’s statistical value (Fesl et al., 2008):

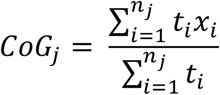

with *X* being the coordinate of voxel *i* in cluster *j* with size *n* and *t* being the statistical value of that voxel.

##### 2.5.4.2. D1 - D5 Distance

Although the distance between D1 and D5 is used to describe the size of the cortical digit representation, it is dependent on exact localization of these digit’s locations. Therefore, this measure was included as a location-based parameter. The Euclidean distance between the CoG of D1 and D5 was calculated using the following formula:

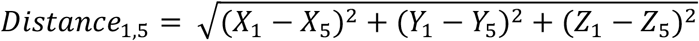

Where *X*_*1*_, *Y*_*1*_, and *Z*_*1*_ and *X*_*5*_, *Y*_*5*_, and *Z*_*5*_ represent the MNI coordinates of D1’s and D5’s CoG, respectively.

##### 2.5.4.3. Volume of Activation

The volume of activation of each digit representation within the anatomically predefined SI ROI was defined as the volume (in mm^3^) of the digit’s activation cluster.

##### 2.5.4.4. Overlap of Activation

The overlap of the activation clusters of two anatomically neighboring fingers (i.e., D1+D2, D2+D3, D3+D4, and D4+D5) was calculated as follows, using the Dice Coefficient (DC) (Dice, 1945):

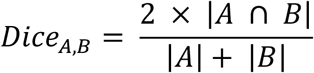

Where |*A ∩ B*| represent the activation volume shared between the two clusters *A* and *B* and |*A*| and |*B*| represent the activation volumes of the clusters *A* and *B*, respectively. It describes the volume of activation that is significantly active for two digits relative to the volume activated by the stimulation of these two digits. Thus, the DC ranges between 0 and 1, with higher values indicating more overlap between the representations of the two digits compared.

#### 2.5.5. Retest Reliability

##### 2.5.5.1. Stability of Activation between Sessions

To investigate the reliability of the activation pattern of both designs, the DC was used in two different ways. First, it was used to test the stability of the single digit activation clusters by calculating the overlap of the clusters belonging to the same digit across the two sessions. High DC values would indicate high stability of that digit cluster, both in terms of location and volume, across the two sessions. The second approach was to investigate the stability of the overlap between neighboring digit activations. For this, the DC was used to calculate the overlap of voxels included in two activation clusters across the two sessions. Again, high DC values would indicate high stability of the overlap of neighboring digit activations in terms of location and size.

##### 2.5.5.2. Correlational Analysis of Map Parameters

To investigate the retest reliability, the map parameters were correlated across the two sessions separately for each mapping design using Pearson correlation. CoG data were correlated separately for each axis (X, Y, and Z), digit, and design, resulting in 30 individual correlation values (3 axes x 5 digits x 2 designs). The D1-D5 distance was correlated separately per design, resulting in two correlation values. The volume of activation was correlated separately for each digit and design, resulting in 10 individual correlation values (5 digits x 2 designs). Lastly, the overlap of activation was correlated separately for each digit-pair and design, resulting in 8 individual correlation values (4 digit-pairs x 2 designs). Thus, in total, 50 correlation values were calculated and tested whether they were significantly larger than zero (one-tailed test). To correct for the multiplicity problem, FDR correction using a linear step-up procedure was used (Benjamini & Hochberg, 1995), values that exceeded a threshold of q(FDR) < .05 were considered significantly larger than zero. Additionally, for each correlation value, the slope and intercept of the best fitting line was calculated using the least squares method. The best fitting line of a highly reliable design is expected to be close to the bisecting angle (i.e., *1x + 0*).

To get a better picture of the individual stability of the CoG locations, the Euclidean distance between each digit’s CoG in the first and second session was calculated.

## 3. RESULTS

Two different somatosensory digit mapping designs, traveling wave (TW) and blocked design (BD), were adapted to clinical conditions and tested for their feasibility of producing single digit BOLD activation maps as well as for their retest reliability in seven neurotypical participants. Data from the first session were used for investigating the feasibility by visual inspection and extracting and comparing parameters concerning the location of the activity as well as the size and overlap of the activation. To investigate the retest reliability of each design, these parameters were also extracted from the second session and correlated to the first session.

### 3.1. Feasibility

#### 3.1.1. General BOLD Activation Pattern

For a general visual impression of the activation patterns elicited by the two designs in the two sessions, significantly activated voxels in volume space were projected onto the flattened cortical surface reconstruction. The mapping designs achieved significant BOLD-activation associated with the passive tactile stimulation of all five digits in all seven participants (*Fig. 3*). This illustrates that both designs have enough power to map all digits’ representations in SI under circumstances mimicking a clinical setting, i.e., limiting the measurement time without compromising on the necessary high spatial resolution. For all participants, the expected anatomical order of the digit activations can be seen along the posterior wall of the central sulcus (dark gray) and the shoulder of the postcentral gyrus (light gray). Activations associated with stimulation of D5 are located most medial and of D1 most lateral, with the remaining digits’ representations positioned in anatomical order between these two endpoints. This sequence of digit activations as well as their location is comparable in all participants, across both mapping designs, as well as across sessions. Also, the area of the elicited BOLD-activations is comparable across the two designs and across the two retest sessions, although with two individual exceptions. One case (*Fig. 3*; first row) exhibits reduced activation during TW compared to BD, in both sessions, but specifically noticeable during the second session. The other case (*Fig. 3*, fourth row) is displaying a distinctly reduced activation in the TW design in the first session only, so this can be generally attributed neither to the TW design, nor to the first session.

**Figure 3.**
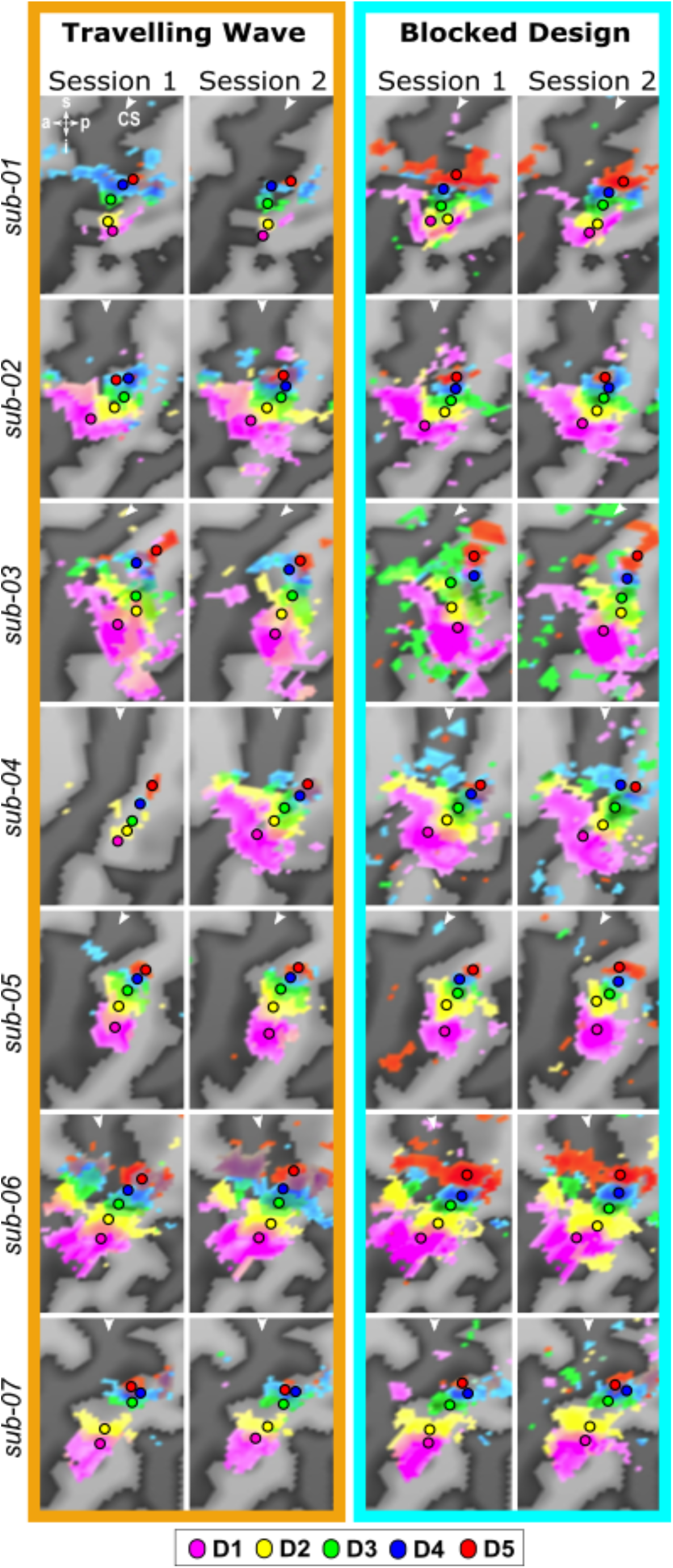
Individual surface-based activation patterns across designs and sessions. Each row depicts the BOLD-activation in response to vibrotactile stimulation projected to the flattened surface of one participant. The left-most dark gray line in each image represents the central sulcus (CS; marked with a white arrowhead) and the light gray line on the right to the CS represents the postcentral gyrus. The top left inset indicates superior (s), posterior (p), inferior (i), anterior (a) directions. The color code of the activation is identical to that of Fig. 1 (i.e., D1 = magenta; D2 = yellow; D3 = green; D4 = blue; D5 = red). All seven participants show significant activations in response to stimulation of all five digits along the posterior wall of the CS, creating a topographic digit map. The location of activation as well as the extent of activation do not seem to change much across sessions and designs with the exception of two cases (first row and fourth row). Note: The surface representation of the activation was done for visualization only; all further analyses are performed in the 3D volume.

#### 3.1.2. Single Digit Center of Gravity: Location

For a comprehensive description of the location of the BOLD activation, the coordinates of all centers of gravity (CoG) of the first session were extracted in 3D volume space (*Fig. 4A*). The CoG of the single digits for TW and BD exhibit distinct digit succession patterns in 3D space across individuals as can be seen in the axial (anterior-posterior and medial-lateral) and sagittal plane (superior-inferior and anterior-posterior). Despite between-subject variations in the position of the succession in 3D space, the group averages of the single digit CoG display the common D1 through D5 succession of distinct digit activation in each coordinate direction.

**Figure 4.**
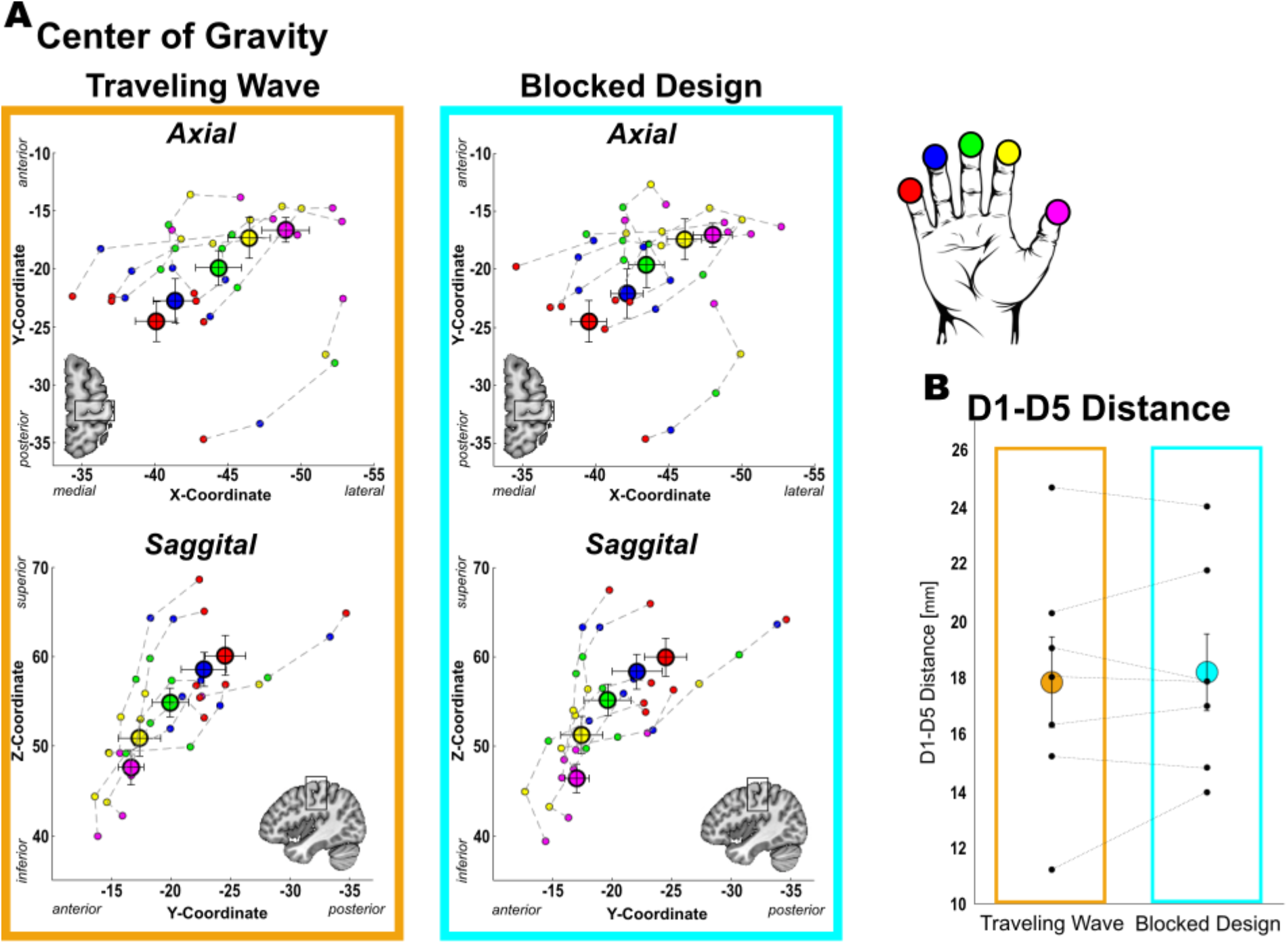
Location based parameter extracted from the digit maps in MNI volume space. **A**. Average locations of each digit’s center of gravity (CoG) in 3D space (large circles) and individual CoGs (smaller circles; each connected succession from D1 to D5 represents one participant). Average and individual CoGs follow the known lateral-to-medial (upper panels) and inferior-to-superior (lower panels) succession. No major differences can be observed between the TW and BD. **B**. Average (large, colored circles) and individual (small, black circles) D1-D5 distances for both designs. Connected individual points belong to the same participant. Again, no major differences can be observed between the two designs. *Error bars represent the standard error of the mean*.

The comparison of the average CoGs of each digit activation across the two mapping designs (*Fig. 4A*) accordingly shows only small differences in location between TW and the BD (Euclidean distance: M = 0.9 mm; Range = 0.5 – 1.6 mm across digits). However, when considering the individual subject level, larger deviations of the individual CoGs between the two mapping designs can be observed (Euclidean distance: M = 2.1 mm; Range = 0.5 – 6.3 mm across digits).

#### 3.1.3. Distance D1/D5 Center of Gravity

The Euclidean distance between the 3D CoG coordinates of D1 and D5 is a measure to describe the extent of the somatosensory hand area (*Fig. 4B*). The within-subject differences of the Euclidean distances between the two mapping designs are, with a mean of 1.0 mm and a range of 0.2 – 2.7 mm across the subjects, considerably small. This is contrasted by the relatively large variation of the total distances between subjects, being in the range of 11.2 and 24.7 mm, which is probably based on the individual differences in the structural layout of the somatosensory digit area, rather than on differences in functional aspects of digit usage.

#### 3.1.4. Single Digit: Area of Activation

The size of a digit’s elicited BOLD-activation was calculated based on the volume of the activation cluster (*Fig. 5A*). On average, the size of the area differs between the digits, with D1 having the largest area of activation (TW: 506.1 ± 179.5 mm^3^; BD: 611.6 ± 121.1 mm^3^). D5 and D4 display the smallest area of activation for TW (D5: 119.0 ± 38.9 mm^3^) and BD (D4: 157.9 ± 28.8 mm^3^), respectively. The general size of the BOLD-activation does vary between participants, some generally larger, some generally smaller, modulated with the digit specific variations. The average sizes of the digit activations are nevertheless similar across the two mapping approaches.

**Figure 5.**
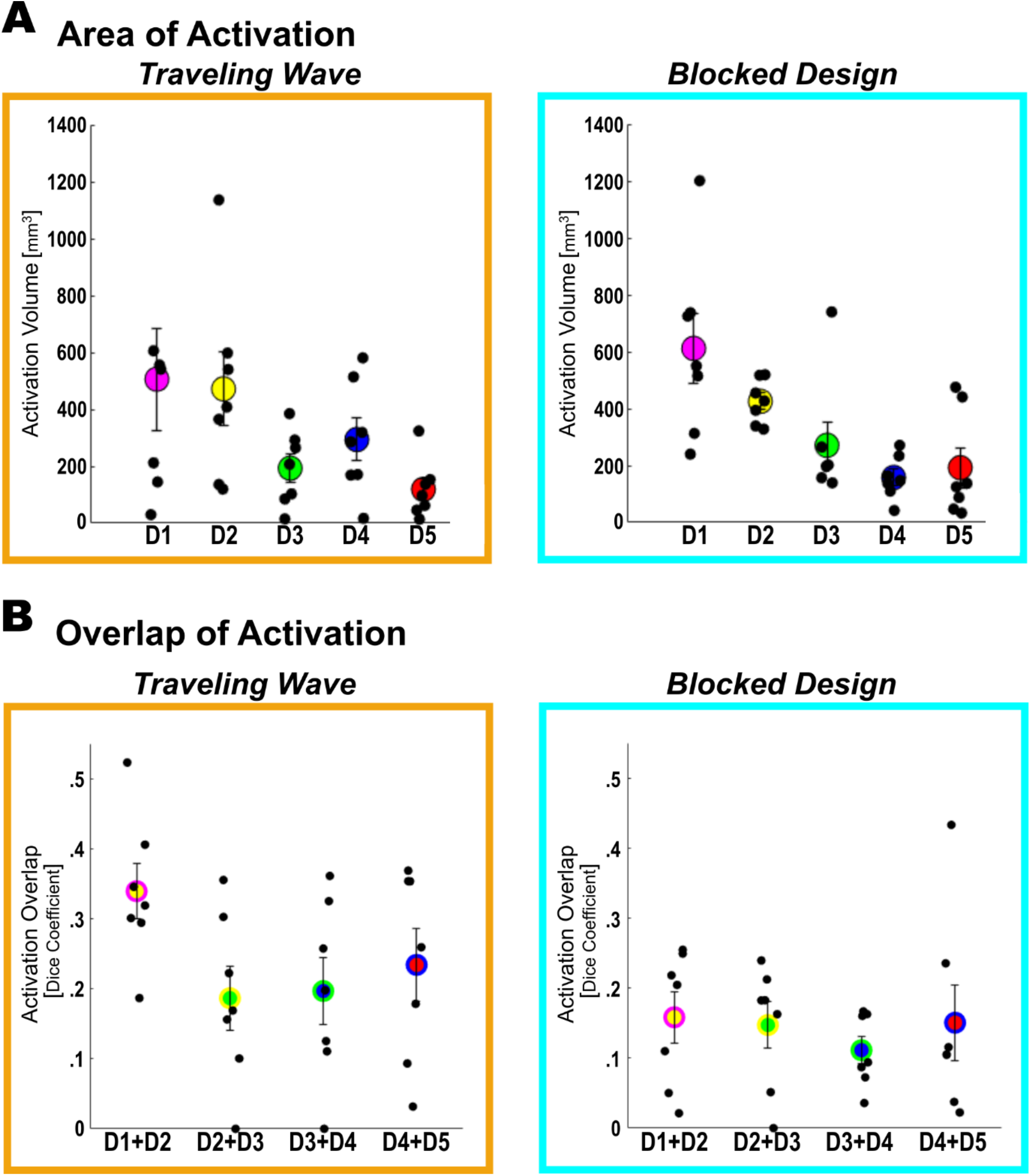
Activation size and overlap of neighboring activation clusters. **A**. Average (large colored circles) and individual (small black circles) activation cluster sizes. D1 and D2 generally occupy the most space while the other three digits (D3, D4, and D5) occupy less space. This general pattern can be observed both for TW and BD. **B**. Average (large two-colored circles) and individual (small black circles) Dice Coefficients (DC) measuring the activation overlap of neighboring digit-pairs. On average, DCs are higher for the TW design as compared to the BD.

#### 3.1.5. Neighboring Digits: Overlap of Activation

The overlap of activation between neighboring digits is quantified by the Dice coefficient (DC), which in this case sets the overlap in relation to the size of the activation clusters of the two neighboring digits, with higher values indicating larger overlap. The comparison of the two mapping designs (*Fig. 5B*) reveals larger average DC values in the TW (Minimum: D2+D3 = .19 ± .05; Maximum: D1+D2 = .34 ± .04) compared to the BD approach (Minimum: D3+D4 = .11 ± .02; Maximum: D1+D2 = .16 ± .04), which suggests less specificity of the single digit activation in the TW approach but is probably a consequence of the difference in signal contrasting within the two analyses. The larger range of DC values for the overlap in all four digit-pairs in the TW design is also evident on the individual level.

### 3.2. Retest reliability across first and second session

#### 3.2.1. Stability of activation

For the retest reliability of the single digit activations, the DC is applied to provide a general description of similarity. The determined overlap of the same digit activation of the first and the second session captures both the potential change in size and location of the activation across sessions. In the DC matrix (*Fig. 6A*), this is represented on the diagonal showing similar average DC values for the five digits in both mapping approaches (TW: .51 – .64; BD: .53 – .75) indicating a 51 - 75% overlap of the single digit activation clusters in the two sessions. The DC values off the diagonal indicate the stability (between the sessions) of the overlap (between each of the four neighboring digit-pairs), by comparing the neighboring digit activation overlap of the first, with the neighboring digit activation overlap of the second session. Again, values are similar between mapping designs (TW: .34 - .43; BD: .19 - .48), however they are generally lower as the values on the diagonal, indicating that the activation overlap between neighboring digits is more variable across sessions compared to the overlap of the single digit activation clusters across sessions. To be able to discriminate the differential effect of location and size of activation on the retest reliability, separate analyses focusing on the degree of variation in these two parameters between the two sessions were undertaken.

**Figure 6.**
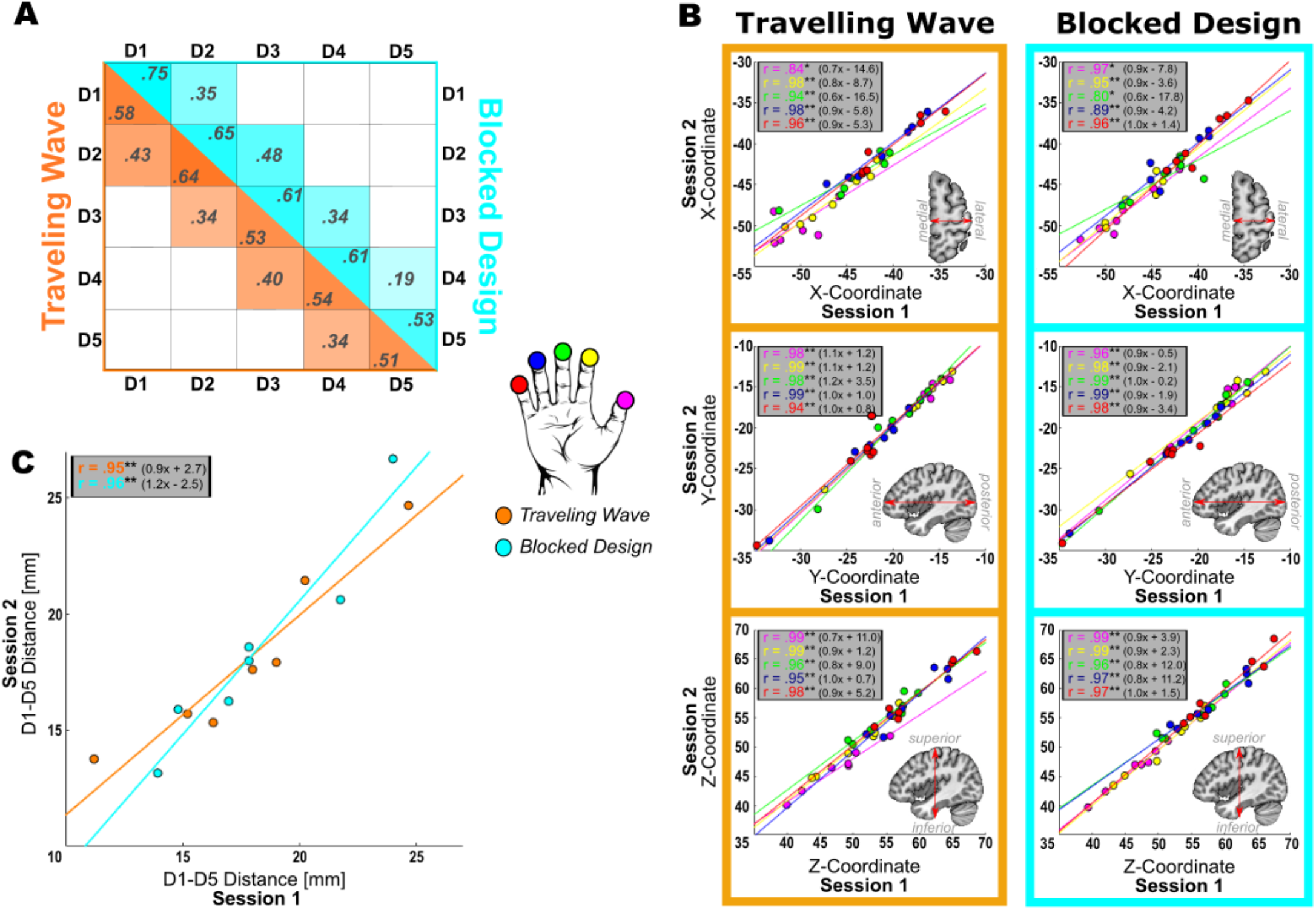
Retest reliability of location-based parameters. **A**. Average Dice Coefficients (DC) describing the overlap of the activation cluster in the first session with the activation cluster of the second session, with higher values indicating more and lower values indicating less overlap. Values on the diagonal represent same-digit overlap, that is, the activation clusters for the same digit are compared between sessions. Values off the diagonal represent comparisons between the area of overlap of neighboring digits between sessions. That is, the area that is common for the activation cluster of two digits in the first session is compared to the common area of the same two digits in the second session. The orange-colored side represents values for the TW design and the cyan colored side represents values for the BD. **B**. All three axes (X, Y, and Z; rows) of each digit’s center of gravity (CoG) were correlated across the first and second session for both designs (columns). Data points represent individual data. All digits in all dimensions achieved significant correlation values indicating high stability of the location of the CoG. Best fitting line described as *(slope)x + (intercept)*. **C**. D1-D5 distances were correlated between the first and second session for both designs (colors). Data points represent individual data. Both designs achieved significant correlation values, indicating that the D1-D5 distance is a stable measure of the extent of the somatosensory hand representation. Best fitting line described as *(slope)x + (intercept)*.*\* p <* .*05; ** p <* .*01; after FDR correction*.

#### 3.2.2. Correlation of Single Digit Centers of Gravity across Sessions

To quantify the retest reliability of the single digit locations for both mapping approaches, each digits CoG X- (medial-lateral direction), Y- (anterior-posterior), and Z- (superior-inferior) coordinates of the first and the second session were correlated (*Fig. 6B*). The results exhibit very high correlation values for all X-Y-, and Z-coordinates, which are significantly larger than zero. Most correlation coefficients even surpass the threshold of q(FDR) < 0.01, with two exceptions for the X-axis of D1 in the TW design (r = .84) and for D3 in the BD (r = .80), which are only significant at q(FDR) < 0.05. Additionally, the best fitting lines are close to the bisecting angle with slopes close to 1 (TW = Mean: 0.91, Range: 0.6 – 1.2; BD = Mean: 0.95, Range: 0.6 – 1.0) and low intercepts (TW = Mean: 5.71, Range: 0.7 – 16.5; BD = Mean: 4.92, Range: 0.2 – 12.0). Together, these high correlation values for all digits and in all three dimensions and best fitting lines indicate a very good retest reliability of the location of the single digit CoG for both designs.

#### 3.2.2. Absolute Distances of Single Digit Centers of Gravity across Sessions

To quantify the differences in CoG locations across sessions, Euclidean distances were calculated between the CoGs of the digits in the first session and the second session for all participants (*Table 1*). Generally, the Euclidean distances confirmed the results of the correlation analysis that the CoG were at similar positions in the two sessions. The average Euclidean distances across participants and digits in TW (M = 1.95, SD = 1.56) and BD (M = 1.58, SD = 1.08) were not significantly different (t(34) = 1.25, p = .22), indicating that there was no difference in retest reliability of the CoG location between the two mapping designs.

**Table 1.** Euclidean distances (in mm) between the CoGs for the same digit in session 1 and session 2 for both TW (gray) and BD (white). Values above 2 mm (i.e., size of one functional voxel) are marked in bold.

|  | D1 |  | D2 |  | D3 |  | D4 |  | D5 |  |
| --- | --- | --- | --- | --- | --- | --- | --- | --- | --- | --- |
|  | TW | BD | TW | BD | TW | BD | TW | BD | TW | BD |
| sub-01 | **4.3** | **2.6** | 0.8 | **2.7** | 1.8 | 0.5 | **3.1** | **2.6** | **5.5** | **2.8** |
| sub-02 | 1.3 | 1.4 | 1.2 | 0.8 | 1.9 | 1.1 | 0.9 | 1.3 | 1.6 | 1.8 |
| sub-03 | **3.9** | 1.6 | 1.4 | 2.0 | 0.7 | **5.2** | **2.8** | 0.9 | **4.9** | **2.7** |
| sub-04 | **5.9** | 0.8 | 1.7 | 1.7 | **4.9** | 1.0 | **2.7** | **4.1** | 0.6 | 0.7 |
| sub-05 | 1.6 | 0.4 | 0.9 | 1.9 | 0.7 | 1.0 | 0.9 | 0.6 | 0.5 | **2.3** |
| sub-06 | 0.7 | 0.7 | 1.8 | 1.0 | **2.6** | 1.2 | 0.5 | 0.6 | **2.7** | 0.3 |
| sub-07 | 0.9 | 1.3 | 1.1 | 1.4 | 0.5 | **2.7** | 0.8 | 0.9 | 0.6 | 0.5 |
| mean<br>± s.e.m. | **2.6**<br>± 0.6 | 1.3<br>± 0.2 | 1.2<br>± 0.1 | 1.6<br>± 0.2 | 1.9<br>± 0.5 | 1.8<br>± 0.6 | 1.7<br>± 0.4 | 1.6<br>± 0.5 | **2.3**<br>± 0.7 | 1.6<br>± 0.4 |

On the individual level, only a few distances were greater than the size of one functional voxel (i.e., 2 mm; marked in bold in *Table 1*; TW = 11 out of 35; BD = 9 out of 35), indicating that the CoG is a reliable measure also on the level of single subjects.

#### 3.2.3. Correlation of D1-D5 Distance across Sessions

The retest reliability of the extent of the somatosensory hand area, estimated by the correlation of the first and second session Euclidean distance between D1 and D5 CoG, also shows very strong correlation values for both mapping approaches (TW: r =.95; BD: r = .96) (*Fig. 6C*). The best fitting lines for both designs are very close to the bisecting angle, showing that the results of the measurements of the D1-D5 distance were very stable across sessions.

#### 3.2.4. Correlation of Single Digit Area of Activation across Sessions

The analysis of the retest reliability of the size of each digit’s cortical activation across sessions revealed considerable variation in the correlation values of each digit’s size of activation between sessions (TW: r = .55 - .78; BD: r = .06 - .97) (*Fig. 7A*). In the TW approach, the area of activation of two digits, D2 and D5, was significantly correlated between sessions. In the BD, three digits, D1, D4, D5, reached significance, while the other two digits (D2 and D3) displayed quite low correlations (D2: r = .06 and D3: r = .14). Also, the best fitting lines do not resemble the bisecting angle in all cases. Especially D1, D2, and D3 as well as D2 and D3 diverge drastically for TW and BD, respectively. These results indicate that the size of digit activations was not always stable across sessions for both mapping designs, as well as that there might be a difference in terms of reliability for the area of activation of different digits.

**Figure 7.**
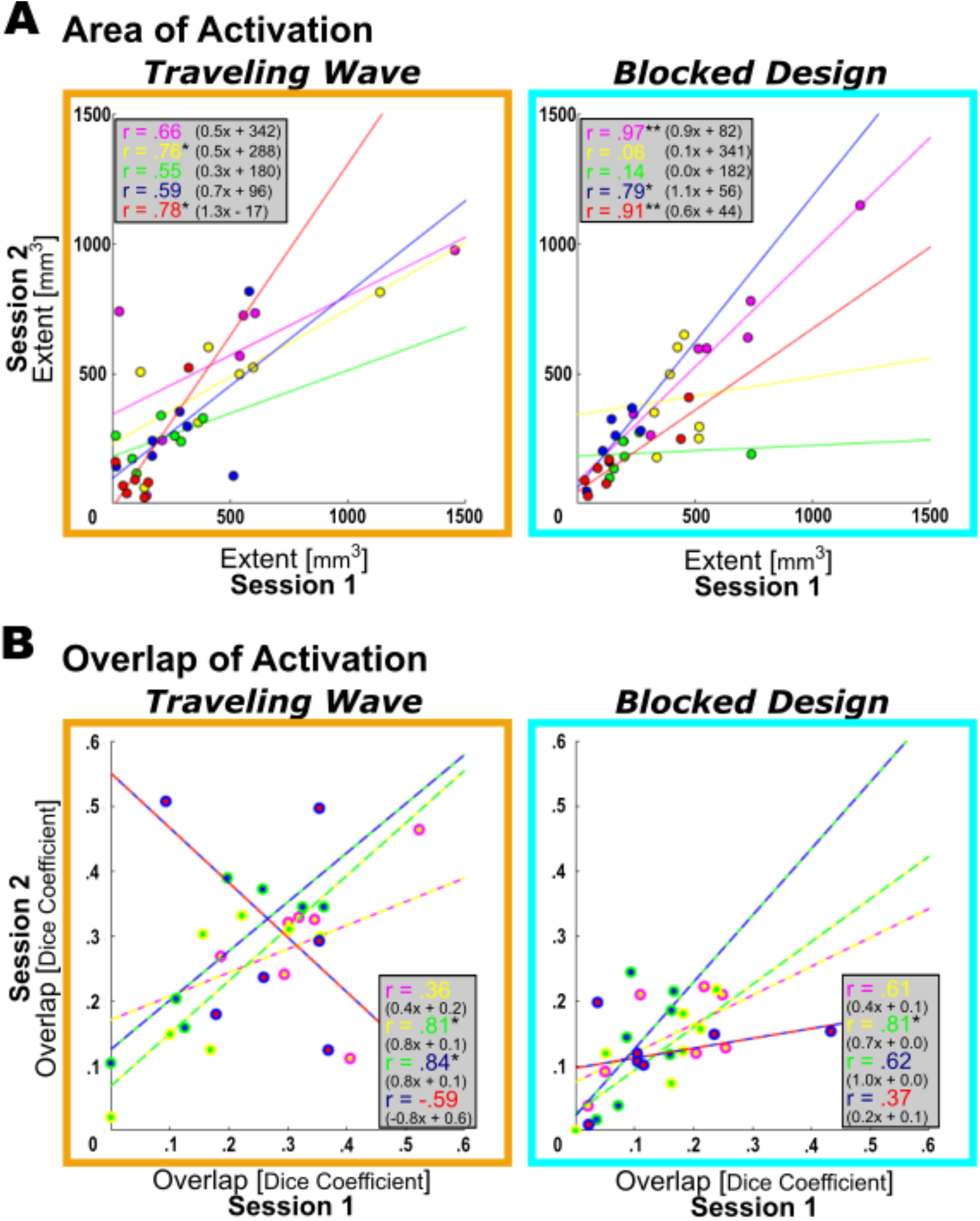
Retest reliability of the size of the activation cluster and activation overlap. **A**. The area of the activation cluster from the first session was correlated to the second session for all five digits (color code identical to previous figures; D1 = magenta; D2 = yellow; D3 = green; D4 = blue; D5 = red). Data points represent individual participants. With a few exceptions, area of activation displayed medium to low correlation values indicating little stability across sessions. **B**. The overlap of the activation of neighboring digits as quantified by the Dice coefficient was correlated between the first and second session for all four digit pairs (D1+D2 = magenta/yellow; D2+D3 = yellow/green; D3+D4 = green/blue; D4+D5 = blue/red). Most correlation values are low and non-significant, indicating little to no stability of the activation overlap across sessions. Best fitting line described as *(slope)x + (intercept). * p <* .*05; ** p <* .*01; after FDR correction*.

#### 3.2.5. Correlation of Neighboring Digits’ Overlap of Activation across Sessions

To assess the retest reliability of the overlap of neighboring digits, the DCs of the activation overlap of the four digit-pairs D1+D2, D2+D3, D3+D4, and D4+D5 were extracted for session 1 and 2 and then correlated (*Fig. 7B*). For the TW, the DC values show a large range of values, which for only two digit-pairs resulted in significant correlations (D2+D3: r = .81; D3+D4: r = .84), one digit pair even produced a negative correlation (D4+D5: r = -.59). For the BD approach, the DCs are less scattered, but the correlations are low except for the D2+D3 digit-pair (r = .81) which was significantly larger than zero. Again, in some cases, the best fitting lines do not resemble the bisecting angle, especially for the digit pairs D1+D2 and D4+D5 for both designs. These results suggest that the size of the activation overlap of most neighboring digit-pairs is not stable across sessions.

## 4. DISCUSSION

This preclinical translational fMRI study investigates the feasibility and retest reliability of travelling wave and blocked design digit mapping approaches adapted for clinical applications. Results confirm that complete and comprehensive digit maps can be obtained despite the constraints for both designs. The locations of the single digit activation CoGs display the commonly observed D5-D1 superior-inferior, medial-lateral and posterior-anterior succession along the central sulcus with high similarity between the designs. Area of digit activation varies between digits but is similar across designs, except for the larger overlap between neighboring digit activation for the TW compared to the BD design. The descriptions of the clinically relevant retest reliability show very high stability of the location of the digit activations between sessions for both designs, revealing that the location of digit representation can be reliably determined. This is contrasted by the area of activation and the overlap with neighboring digits activation exhibiting a considerable variability and only moderate stability across sessions making it a less reliable descriptor of the digit activation. In the following these results are assessed in relation to existing data and future directions for optimization are discussed.

### 4.1. Feasibility of Somatosensory Digit Mapping in a Clinical Adaptation

Both mapping designs, TW and BD, yielded significantly activated voxels in response to passive tactile stimulation of all five digits in both sessions for all participants, demonstrating the general feasibility of the adapted procedures to map the entire cortical hand representation in SI.

The considerably reduced spatial resolution of 2 mm isotropic (8 mm^3^), compared to the 1.5 mm isotropic (3.375 mm^3^) at 3T or even 1.2 mm isotropic (1.7 mm^3^) at 7T used in basic research studies, does limit the spatial specificity of the signal, and increases partial volume effects, considering the narrow width (< 2 mm) of SI (Fischl & Dale, 2000; J. R. Meyer et al., 1996). However, the findings that activation for all 5 digits was present in all participants shows that this disadvantage is counterbalanced by the larger voxel volume yielding a stronger signal and lower sensitivity to motion artefacts, which turns into an important benefit for the implementation in a clinical population.

The analysis of the location and area parameters of the single digit activation provide a valid description of the topography that is in line with the results from basic research studies: the known succession of the single digit centers of activation along the medial-lateral, superior-inferior and posterior-anterior trajectory of the posterior wall of the central sulcus (Besle et al., 2013; Kurth et al., 2000; Martuzzi et al., 2014; Nelson & Chen, 2008; Sanchez-Panchuelo et al., 2010), the comparable D1-D5 Euclidean distances gauging the somatosensory hand area (Härtner et al., 2021; Schweisfurth et al., 2018), and the digit specific sizes of activation elicited by the tactile stimulation (Martuzzi et al., 2014; Schweisfurth et al., 2018) confirm the viability of the adaptations.

In addition to the general feasibility of the clinically adapted approaches in obtaining valid digit maps, it is important to emphasize that both mapping designs, TW and BD, yielded comparable results in almost all location- and area-based measures, despite the differences in stimulation and analysis, which speaks to their robustness. Furthermore, this may confirm a number of decisions made concerning the analysis and parameterization. First, the structural definition of BA 3b as the region of interest, although activation from BA3a or, more plausible, activation from BA1 cannot be excluded. Second, the removal of voxels significantly activated by three or more digits, potentially representing BOLD-activation originating from blood vessels and not gray matter, which could lead to false interpretation of the fMRI-based digit map under the specific conditions of lower magnetic fields at 3T and lower spatial resolution (Besle et al., 2014; Nelson & Chen, 2008; Schweizer et al., 2008). Third, the limitation to one cluster per digit, following the description of the single digit representation within BA3b obtained with electrophysiology in monkeys (Iwamura et al., 1983; Merzenich et al., 1978; Recanzone et al., 1992). Fourth, the selection of the center of gravity within this cluster to represent the location of the digit representation (Vidyasagar & Parkes, 2011; Weibull et al., 2008). Although, more studies and analyses, especially with patient populations, are needed to put these decisions further on trial.

The center of gravity approach was implemented in an automatized approach to extract each digit’s activation cluster based on its size, the presence of the peak voxel, and in the case of diverging choices, also its location. This expert-based pipeline is, again, an adaptation to the clinical setting, with its strong need for automatized procedures, reducing the burden on health care practitioners, especially for rare procedures as this detailed fMRI analysis. A recent approach to create a standardized and automatized analysis pipeline for somatosensory digit mapping data based on the high spatial resolution fMRI and additional MR-angiography information (Pfannmöller et al., 2016) was informative, but not applicable for the clinical application due to its high spatial resolution and the need for an additional MR-angiography scan.

Though not formally confirmed, a noticeable difference between the two mapping designs is the larger and more variable overlap of the activation of neighboring digits in the TW compared to the BD. This, however, can most probably be ascribed to the different analysis approaches applied to the different stimulation schemes. The BD was analyzed with the standard GLM, and each digit predictor was contrasted against the average of the predictors for the remaining digits. For the phase-encoding cross-correlation analysis applied to the TW stimulation no contrast was applied. Moreover, the inclusion of two models (see Methods and Materials) for the cross-correlational analysis of each digit could have contributed to the increased activation overlap between neighboring fingers in the TW design. Differences in the adaptable FDR thresholding also have to be considered, as the level of thresholding influences the percentage of voxels being activated by two digits and consequently the extent of the overlap (Besle et al., 2014). The specific analyses for this clinical adaptation were chosen to closely follow the approaches of published studies (Kolasinski, Makin, Jbabdi, et al., 2016; Schweisfurth et al., 2018) to allow the comparison of the results to the results of these basic research studies.

Taken together the comparison of the TW and BD mapping results show each design’s power and capacity to map the somatosensory digit area even when taking into account possible limiting factors of a clinical setting. The two approaches achieve comparable results for the location of the single digit activation as well as for the volume of the activated area, which are analogous to basic research studies having the advantage of higher spatial resolution and higher field strengths.

### 4.2. Retest Reliability

The Dice coefficients, describing retest reliability by comparing the overlap of the single digit activation of the first and the second session, resulted in 51% – 75% correspondence of the activation for both mapping designs. These values are in the same range as reported by Kolasinski and colleagues (Kolasinski, Makin, Jbabdi, et al., 2016) for their intra-subject single digit map reproducibility of an active mapping task applying the TW design. Despite the encouraging correspondence of this clinical adaptation with the results of the ultra-high field / high spatial resolution study, the actual values of the similarity between measurements are much lower than hoped for. A clinical implementation of the digit mapping, with the prospect of detecting changes associated with recovery, as well as with therapeutic interventions, requires a higher within-subject retest reliability. Since the Dice coefficient, as a similarity measure, takes into account the location as well as the area of the digit activation, separate analyses of the retest reliability of the location and of the areas of the single digit activations were conducted to investigate the contribution of each of the components.

#### 4.2.1. Retest Reliability of Location Based Measures

The subsequent correlational analyses of the single digit CoGs between the two sessions show high values with the correlation being very close to the bifurcation line for both designs, indicating a high retest reliability of the location of the single digit CoGs for all participants and both designs. These results converge with previous reports of the circumscribed across-session variability of the location of digit activations. Values of the between session variance for somatosensory fMRI measurements at 3T are not available, since location stability across sessions was, especially in the early digit mapping studies in humans, not explicitly investigated. This changed with ultra-high field 7T fMRI, providing higher spatial resolution to investigate not only reorganization, but potentially also digit map adaptations. There, data associated with retest reliability are occasionally reported as part of the general descriptions, showing a highly stable location of digit activation comparing different runs within the same session (Stringer et al., 2011) as well as between sessions with longer intermediated time periods (Martuzzi et al., 2014). The actual reported average values of the distances between sessions are in the same range as the values in the present study (Martuzzi et al. 2014: min = 1.7 ± 1.2 mm D1; max = 3.2 ± 3.1 mm for D2 | present study: min = 1.2 ± 0.1 mm D2; max = 2.6 ± 0.6 mm for D1) both demonstrating a displacement of an equivalent of 1-2 functional voxels. A confirmative retest reliability is reported in a 9.4T, high spatial resolution MRI study in squirrel monkeys, assessing the limits for the description of reorganization/adaptation in BA3b across measurements weeks to month apart. The locations of the single digit activation peak voxels between sessions show an average variation of 0.5 ± 0.15 mm, which considering the in-plane spatial resolution of 625 × 625*μ*m^2^, also correspond to a displacement of the size of 1-2 functional voxels (Zhang et al., 2010). This variability of the single digit location across sessions by on average 1-2 voxels, even across different magnitudes of magnetic fields and spatial resolutions, demonstrates a proficient stability and retest reliability of the location of single digit activation, which is promisingly matched in the present clinical implementation. Although, it has to be pointed out that, on the individual level, in 43% of the participants (3 out of 7), 40% of the digits (2 out of 5) showed a displacement of the CoG of three voxels (4.1 - 5.9 mm), which is due to the relatively course spatial resolution, already within the range of the distance to the CoG of the neighboring digit.

#### 4.2.2 Retest Reliability of Volume Based Measures

Contrary to the location, the area of activation and the overlap between the areas of neighboring digit representations shows only medium to low correlation values and medium to large deviations from the bisecting line. This indicates more pronounced variations in the elicited areas of the single digit activation across sessions and consequently a lower retest reliability.

Considerable within-subject variations of the extent of BOLD activations between sessions, despite using identical stimulation and measurement regimes, are not unusual and have been investigated and described in detail (e.g., McGonigle et al., 2000) providing the basis for the development of suitable preprocessing, analysis, and thresholding strategies (e.g., Smith et al., 2005). Concerning the specific topic of the extent of single digit activation across sessions, actual descriptions are, again, sparse and reveal a variety of sources. Martuzzi and colleagues (2014) report a general decrease in the volume across all single digit activations in the second sessions, which the authors comprehensively explain by a drop in attention. In the present study, a counting task was associated with the tactile stimulation to assure a constant focus of attention on the stimulation across the sessions, and to prevent a general loss of signal across all single digit activations in the second session. A more general approach was taken by the already referred to ultra-high field fMRI study in squirrel monkeys, in which the large variations in the size of the digit activation across sessions are attributed to changes in the signal to noise ratio. The comparison of different thresholding procedures revealed that a flexible, and stricter, thresholding approach resulted in much smaller areas of activation, which were more reproduceable across sessions than the larger activations resulting from a fixed or a combined fixed/flexible thresholding approach (Zhang et al., 2010). The present study applies the False Discovery Rate (FDR) (Benjamini & Hochberg, 1995) as a flexible thresholding approach, which adapts to the general difference of BOLD signal between sessions, partially counterbalancing its effect on the number of activated voxels. But even under the condition of FDR threshold adapting from measurement to measurement, there is a considerable difference in area variability between the single digits causing digit specific decrements in the retest reliability. One of the possibilities to reduce this variability is to increase the threshold (see Zhang et al., 2010) to select voxels with strong enough activation that the BOLD signal fluctuations lose relevance (McGonigle et al., 2000). But this approach, although very successful in increasing the retest reliability of the area of BOLD-activation, would in the extreme case approximate the area around the peak voxel and thereby converge with the location retest reliability. More strict thresholds would also reduce the specificity for the potential and probably subtle changes in the extent of the activation reflecting changes in the underlying neuronal digit representation, as would clinically be expected associated the adaptation within the digit map in response to recovery and/or treatment interventions changes in the digits and hand usage. Further studies are needed to balance out this conflict between reliability and specificity, both being important for the description of the layout of the digit map as well as for the detection of changes within this layout.

#### 4.2.3 Retest Reliability of Overlap Based Measures

The retest reliability of the overlap of neighboring digit activation shows an, at least equally large, variation and low correlation across the two sessions as the retest reliability of the area of activation. This is also reflected in the low Dice coefficients showing an average correspondence of only 36% between the neighboring digit overlaps in the two sessions, also indicating larger fluctuations of the overlap of neighboring digit activation. The only other study mentioning the retest reliability of the overlap of digit activation across sessions depicts the within subject fluctuation of Dice coefficients of the digit activation overlap between sessions and show that the between subject variation of the overlap between digits and sessions is larger than the within subject variation (Kolasinski, Makin, Jbabdi, et al., 2016). Enhanced intra-subject reliability could be obtained, as discussed with the area of activation, by more conservative thresholds, reducing the area of activation, and consequently reducing the overlap between neighboring digit activations. Again, additional high spatial resolution fMRI studies at ultra-high fields are needed to clarify the different factors influencing the reported overlap of activation of neighboring digits (Besle et al., 2014; Kolasinski, Makin, Jbabdi, et al., 2016; Schweizer et al., 2008) as well as addressing the general question of how the layout of the individual neuronal digit representations can reliably be captured, to provide a basis for an adaptation into the specifics of clinically applicable mapping procedures.

### 4.5. Limitations of the study

Despite the clinical focus of this implementation of digit mapping procedures, this research was not performed with patients but with young neurotypical participants. This limitation to the general validity of the approach had to be taken into account for two reasons, namely to be able to adequately describe the feasibility as well as the retest reliability of the clinical adaptation. To specify the feasibility, two different mapping approaches were compared. Consequently, multiple measurements had to be performed within a session, which would not have been reasonable nor acceptable for older adults or patients. The description of retest reliability on the other hand, required two of these multiple measurement sessions with the additional prerequisite that no alterations in the topographical organization of the digit map are expected to occur during the intermediate time, which would be an unrealistic prospect in patients. There are, evidently, clear limitations in the transfer of the results of the clinical adaptation of the procedures, if they are tested in neurotypical young adults only. The generalization of findings from neurotypical participants to patient populations must be approached with caution, since even fundamental parameters can differ, such as a slightly altered hemodynamic response function in stroke patients (Bonakdarpour et al., 2007), calling for an adapted fMRI analysis in these patients.

Likewise, the sample size of seven participants is rather small, though comparable to other studies investigating the reliability of somatosensory digit mapping procedures (Kolasinski, Makin, Jbabdi, et al., 2016 (n = 9); Martuzzi et al., 2014 (n = 10); Stringer et al., 2011 (n = 6)). That sample size does not always matter, can be read in the insightful neuroimaging analysis study of Marek and colleagues (2022) establishing that big samples of thousands of individuals can be necessary to perform specific studies, e.g., brain wide associations, but the authors also do not hesitate to stress the importance of small sample within-subject studies for clinical care. Moreover, as the aim of the current study was to investigate the feasibility and reliability of somatosensory mapping for a clinical context, the focus was on the individual, rather than the group level. When the obtained digit map parameters are expected to be used for diagnostics or prognosis of treatment outcome, the description of the variability on the level of the individual has to be included in the description of the reliability.

## 5. CONCLUSION

This study shows that the fMRI-based mapping of the somatosensory digit representations in SI can, even on the individual level, be feasible, when taking certain limitations of the clinical setting into consideration. The valid description of the general topography within BA3b of the somatosensory cortex exhibiting the succession of the single digit representation is an important prerequisite to determine potential reorganization of the digit maps in clinical conditions. The additional description of the retest reliability, showing a high stability of the location of the digit representation over time, compared to a lower replicability of the extent of the digit representation and its overlap with neighboring digit representations. While no major differences in terms of feasibility nor retest reliability were found between the two mapping designs, the more conventional statistical analysis procedure might favor the blocked design for clinical application. This is a promising first step towards the clinical assessment of somatosensory cortex reorganization due to trauma and digit map adaptability due to interventions.

## 7. Acknowledgements and Funding

We mourn the untimely and irreplaceable loss of our friend and colleague, Amanda Kaas. Her enthusiasm, inclusiveness, and persistence made this study possible.

This work was supported by the NWO Research Talent Grant [406.18.565] to T.S.; Leibniz ScienceCampus Primate Cognition Outgoing Grant [LSC_OG2016_01] and Seed Fund [LSC_SF2018_06] to R.S. Financial support for data acquisition was provided by FPN-MBIC (Maastricht University, the Netherlands) funding to A.K.

## 8. Author Contributions

**Till Steinbach:** Conceptualization, Methodology, Investigation, Software, Validation, Formal Analysis, Data Curation, Writing – Original Draft, Visualization, Funding Acquisition

**Judith Eck:** Conceptualization, Methodology, Investigation, Software, Formal Analysis, Writing – Review & Editing

**Inge Timmers:** Conceptualization, Methodology, Investigation, Writing – Review & Editing **Emma Biggs:** Conceptualization, Methodology, Investigation, Writing – Review & Editing **Rainer Goebel:** Conceptualization, Resources, Writing – Review & Editing, Supervision, Funding Acquisition

**Renate Schweizer:** Conceptualization, Methodology, Validation, Writing – Original Draft, Writing – Review & Editing, Supervision, Funding Acquisition

**Amanda Kaas:** Conceptualization, Methodology, Software, Validation, Formal Analysis, Investigation, Data Curation, Supervision, Project Administration, Funding Acquisition

